# Gut microbiome carbon and sulfur metabolisms support *Salmonella* during pathogen infection

**DOI:** 10.1101/2024.01.16.575907

**Authors:** Ikaia Leleiwi, Katherine Kokkinias, Yongseok Kim, Maryam Baniasad, Michael Shaffer, Anice Sabag-Daigle, Rebecca A. Daly, Rory M. Flynn, Vicki H. Wysocki, Brian M. M. Ahmer, Mikayla A. Borton, Kelly C. Wrighton

**Affiliations:** Department of Cell and Molecular Biology, The Colorado State University, Fort Collins, CO, USA; Department of Soil and Crop Sciences, The Colorado State University, Fort Collins, CO, USA; Department of Microbiology, Immunology, and Pathology, The Colorado State University, Fort Collins, CO, USA; Department of Chemistry and Biochemistry, The Ohio State University, Columbus, OH, USA; Department of Microbial Infection and Immunity, The Ohio State University, Columbus, OH, USA; Resource for Native Mass Spectrometry Guided Structural Biology, The Ohio State University, Columbus, OH, USA

## Abstract

*Salmonella enterica* serovar Typhimurium is a pervasive enteric pathogen and an ongoing global threat to public health. Ecological studies in the *Salmonella* impacted gut remain underrepresented in the literature, discounting the microbiome mediated interactions that may inform *Salmonella* physiology during colonization and infection. To understand the microbial ecology of *Salmonella* remodeling of the gut microbiome, here we performed multi-omics approaches on fecal microbial communities from untreated and *Salmonella*-infected mice. Reconstructed genomes recruited metatranscriptomic and metabolomic data providing a strain-resolved view of the expressed metabolisms of the microbiome during *Salmonella* infection. This data informed possible *Salmonella* interactions with members of the gut microbiome that were previously uncharacterized. *Salmonella-*induced inflammation significantly reduced the diversity of transcriptionally active members in the gut microbiome, yet increased gene expression was detected for 7 members, with *Luxibacter* and *Ligilactobacillus* being the most active. Metatranscriptomic insights from *Salmonella* and other persistent taxa in the inflamed microbiome further expounded the necessity for oxidative tolerance mechanisms to endure the host inflammatory responses to infection. In the inflamed gut lactate was a key metabolite, with microbiota production and consumption reported amongst transcriptionally active members. We also showed that organic sulfur sources could be converted by gut microbiota to yield inorganic sulfur pools that become oxidized in the inflamed gut, resulting in thiosulfate and tetrathionate that supports *Salmonella* respiration. Advancement of pathobiome understanding beyond inferences from prior amplicon-based approaches can hold promise for infection mitigation, with the active community outlined here offering intriguing organismal and metabolic therapeutic targets.

## Introduction

Non-typhoidal *Salmonella*, including *Salmonella enterica* serovar Typhimurium (hereafter *Salmonella*), cause more than 1.3 million infections and hundreds of deaths annually in the United States (1,2). The global burden of non-typhoidal *Salmonella* is magnitudes larger and the World Health Organization reports 550,000,000 infections globally per annum (3). *Salmonella* is often transmitted by contaminated food products and is harbored in zoonotic reservoirs of common food animals like chickens and pigs (4). Recently, *Salmonella* serovars were responsible for multiple outbreaks linked to alfalfa sprouts, cantaloupe, seafood, and peanut butter (5). Given the global emergence of multidrug-resistant *Salmonella* strains (6–8) there is an urgency to develop alternatives to traditional antibiotic therapy, which could leverage understanding of the metabolic targets that sustain pathogen infection.

Much of what is known about *Salmonella* metabolism during infection stems from work done in gnotobiotic mice, or in murine models that require pre-treatment with antibiotics to facilitate infection (9–13). Such models disrupt understanding of natural microbiota interactions during infection. From these prior studies it was demonstrated that *Salmonella* harnesses host inflammation, creating a range of respiratory electron acceptors including oxygen, oxidized nitrogen, and oxidized sulfur species (11,14,15). It was also shown that *Salmonella* utilizes microbiota-derived carbon sources like succinate, 1,2 propanediol, and ethanolamine (10,16–18). Additionally, prior studies demonstrated that *Salmonella* utilized host-derived lactate and tetrathionate as a carbon source and electron acceptor respectively (9,11). While, these findings laid an important foundation of physiological processes germane to *Salmonella* expansion and persistence in the gut (19–25), there is limited work published on the post-colonization microbiota metabolic interactions with *Salmonella*. Further, these metabolisms have not been collectively examined through community-wide transcriptome-based analyses, and until now there has been no holistic evaluation of *Salmonella* sulfur and carbon metabolism amidst an unperturbed microbiome.

The CBA mouse offers a more ideal model for observing *Salmonella* gastroenteritis effects on the gut microbiome, as it avoids pre-treatment with antibiotics while occluding *Salmonella* systemic infection (26,27). Importantly, this disease model mimics the inflammatory response to *Salmonella* in the human gastrointestinal tract and is becoming a mainstay of *Salmonella* research (10,14,20,28–30). In our prior work with CBA mice we repeatedly showed that *Salmonella* significantly restructured the microbial community of the gut, reducing the relative abundance of many dominant *Clostridia* and *Bacteroidia* while enriching for rare *Bacilli* and selected membership from *Clostridia, Bacteroidia*, and *Verrucomicrobiae* (31,32). Importantly, we created the first metagenome assembled genome (MAG) catalog from the CBA model (CBAJ-DB), sampling over 2,000 MAGs from *Salmonella* infected and non-infected mice (31). The CBAJ-DB revealed unique bacterial lineages from *Bacilli, Bacteroidia,* and *Clostridia* not found in catalogs from other mouse strains and showed that *Salmonella*-impacted microbial communities harbored greater potential for respiration and lower potential for butyrate production (31). While important foundationally to advancing pathobiome research, our prior research was entirely DNA based inference. Overall, this lack of gene expression or chemical data limited our ability to fully expound pertinent *Salmonella* metabolic processes and obscured possible microbial exchanges between *Salmonella* and other gut members.

Here we utilized the CBA genomic resource to contextualize paired metabolome and metatranscriptome data from non-infected and *Salmonella* infected CBA/J mice. Metatranscriptomics offers a more active physiological status of *Salmonella* to reveal important interfaces with the native microbiota. First, we profiled *Salmonella* gene expression during the late stage of infection, providing some of the first insights into microorganisms metabolically active during *Salmonella* infection. Our multi-omics approach interrogated metabolic pathways co-expressed during infection, revealing new microbial contributions to carbon and sulfur cycling and the re-assignment of metabolites historically considered only host derived. Ultimately, multi-omics allowed renewed examination of the ecology during pathogen infection in a more robust community context, providing a microbiome interaction framework for future insights on diseases of the gastrointestinal tract.

## Results and Discussion

### Experimental design targeting the metabolic niche of *Salmonella* in the inflamed gut

Using 16S rRNA amplicon sequencing we temporally evaluated the feces from CBA/J mice infected with 10^9^ colony forming units of *Salmonella* (n=44 mice) and an uninfected cohort (n=23 mice) left without *Salmonella.* From a subset of these mice, we report a collection of metabolite and metatranscriptome samples collected from seven *Salmonella* infected (infected, I1-I11) and seven non-*Salmonella* infected (uninfected, U1-U8) mice (Fig. 1A). Due to limited fecal sample at the later stages of infection, metabolites were collected on day 12 from each of the 14 mice and metatranscriptomes were collected on day 11 from five mice in each treatment (Fig. 1A).

**Fig. 1:**
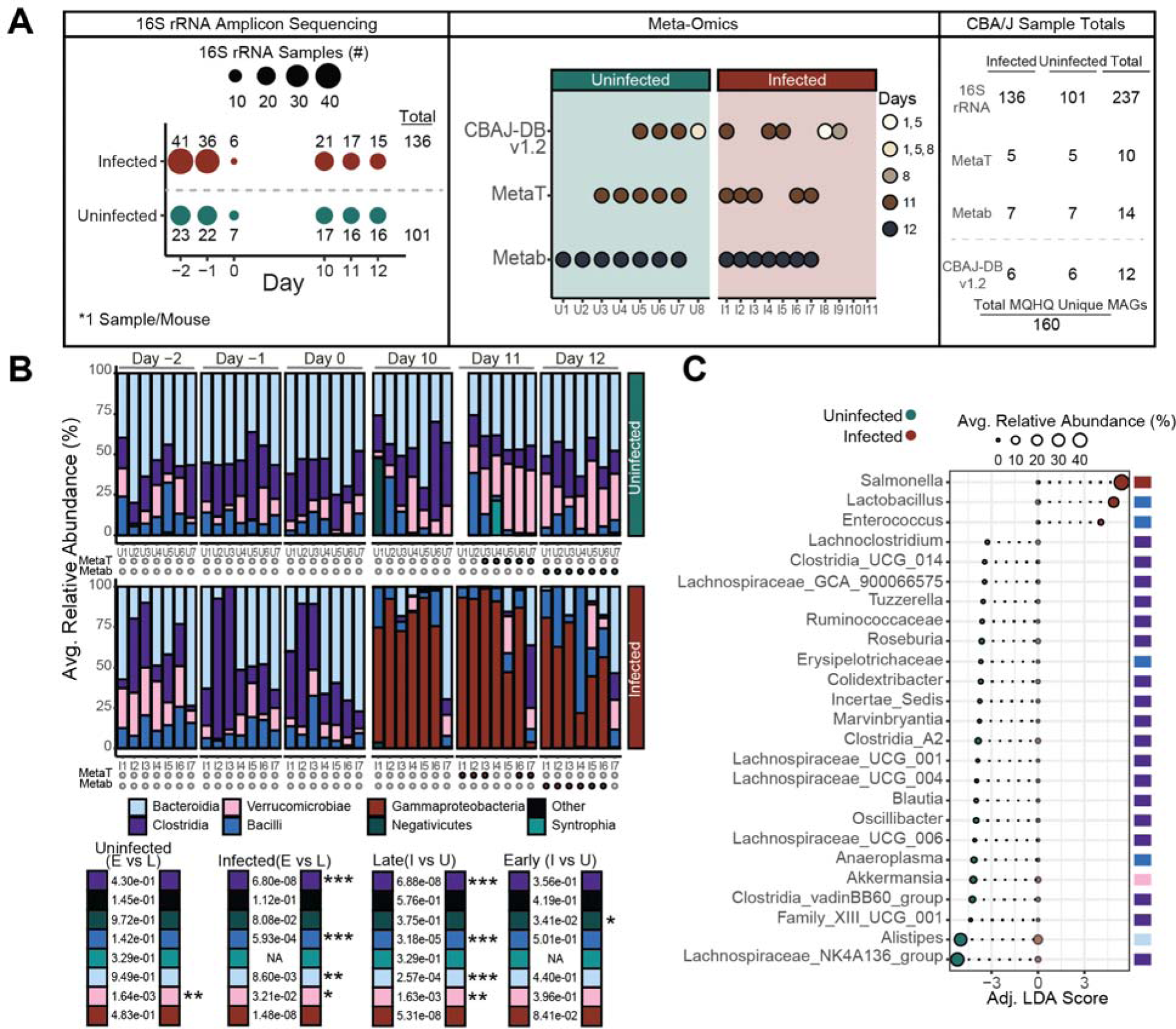
*Salmonella* infection enriches for *Bacilli* in communities and selects for distinct bacterial community membership. **A** Experimental sampling scheme showing total fecal samples taken on each day for 16S rRNA amplicon sequencing (left), and multi-omics analysis (middle). *Salmonella* treatment start is on day 0 and specific sample totals by treatment are listed in the right figure panel. Mice with multi-omics data are designated with a “U” if they were in the uninfected group (n=8) or with an “I” if they were in the *Salmonella* infected group (n=11). **B** Stacked bars showing the ASV class distribution of mice that have metatranscriptomic and metabolomic data. Mouse U1 Day 11 sample was omitted via ASV table filtering (see methods). Significant differences (Wilcoxon Rank Sum) between classes of either early timepoints (E; days −2, −1, and 0), late time points (L; days 10, 11, and 12), infected samples (I), or uninfected samples (U). **C** Linear discriminant analysis of 16S rRNA amplicon data from late timepoint samples (day 10-12) from the subset of mice with metatranscriptomic and metabolomic data. Points are sized by relative abundance of each genus within a treatment (infected or uninfected) and are colored by treatment where points aligning with x-axis value 0 are the relative abundance of each genus in the non-significant treatment. Genera classes are listed by color on the right of the plot.

With 16S rRNA amplicon sequencing performed at days −2, −1, and 0, as well as days 10, 11, and 12 post-infection we verified later stage infection communities on days 10, 11 and 12 were not statistically different from one another and were distinct from the pre-infection and uninfected communities (Fig. 1B). Consistent with two prior studies using the CBA model (31,32), a linear discriminant analysis of the subset of mice used for multi-omics analyses indicated *Lactobacillus* and *Enterococcus* were the predominant *Bacilli* enriched with *Salmonella* (Fig. 1C) (31). In fact, these two members were the only genera that showed positive significant Spearman correlations with *Salmonella* at later timepoints of severely infected mice (*Salmonella* relative abundance ≥ 25% in at least one timepoint) (Fig. S1A).

Genera (n=19) with significant positive correlations to at least one other taxa during infection include *Akkermansia*, *Alistipes*, 14 *Clostridia* genera (Fig. S1A). These results paired with the discriminant analysis results indicate a strong association of *Lactobacillus* and *Enterococcus* with *Salmonella* during later stages of infection and provide evidence of a distinct bacterial community associated with *Salmonella* infection.

Since the earlier publication of CBAJ-DB (v1.0) (31), we have updated the collection to v1.2, by adding metagenome assembled genomes (MAGs) reconstructed from 6 new metagenomes from CBA mouse feces that were either infected or uninfected with *Salmonella*, resulting in 47 new MAGs in the non-redundant medium and high quality database (contamination < 10% and completeness ≥ 50%) (Fig. S1B). The database is publicly available (Doi: 10.5281/zenodo.8395759) and includes 3,667 total MAGs clustering at 99% genome identity to 160 non-redundant, quality MAGs. We confirmed CBAJ-DB v1.2 included representation of *Enterococcus* (n=5 MAGs) and *Lactobacillus* (n=6 MAGs) recovered during infection. We note that the amplicons detected here had perfect matches to MAG recovered 16S rRNA genes from the highest quality *Lactobacillus johnsonii* MAG (Data 1), indicating the co-occurring strains during *Salmonella* late infection were well represented in our genomic catalog.

The CBAJ-DB v1.2 acted as the basis for the analysis of metatranscriptomics reported here. The ten metatranscriptomes sampled on day 11 from uninfected and post-infection mice were mapped to this database (Fig S1B). Metatranscriptomes were sequenced to a max depth of 11.4 Gbps and to a mean depth of 6.8 Gbps (mean 53,689,578 paired 151 bp reads) to ensure read recruitment to non-*Salmonella* and lower abundance members of the microbial community (Fig. S1B). When we rarified the data to 2.5 Gbps, a common value in gut microbiome analyses, it decreased our MAG representation in the infected metatranscriptome by 51%, indicating the value of our deeper sequencing (Data 2, see methods). Seven metagenomes from infected mice sampled on day 8 or day 11 were also mapped to the CBAJ-DB v1.2 (Data 1) to ascertain genome relative abundance in the infected community (Figs. 1A, S1B). The mapped metagenomes were deeply sequenced to a max depth of 68.4 Gbps and a mean depth of 27.4 Gbps (mean 191,655,610 paired 251 bp reads) to avoid sequence saturation from high abundant *Salmonella* and resulting in the recovery of a combined 160 medium and high-quality MAGs all with detectible read mapping from infected metagenomes (Fig. S1B).

### Salmonella expresses genes for diverse carbon and energy metabolic strategies

This to our knowledge is the first *in vivo* metatranscriptomic analysis of *Salmonella* with a native microbiota (Fig. 2A), offering insights into the transcriptionally active microbial strains and their expressed metabolisms during late-stage infection when *Salmonella* relative abundance often exceeded 50% (Fig. 1B). On average the *Salmonella* genome recruited 20% of the community metatranscriptome reads from the five infected metranscriptomes from day 11 (Data 2). It is well documented that *Salmonella* initiates host inflammation that creates oxidized electron acceptors that the pathogen then uses to respire and outcompete the normal flora (11,14,15). It is thought that the respiration provides an energetic advantage over the obligatory fermentative commensal members of the gut microbiota (33). However, it was not known which of these *Salmonella* respiratory metabolisms were expressed during late stages of infection, and if other members of the community were metabolically active, and how they persisted in these more oxidative conditions.

**Fig. 2:**
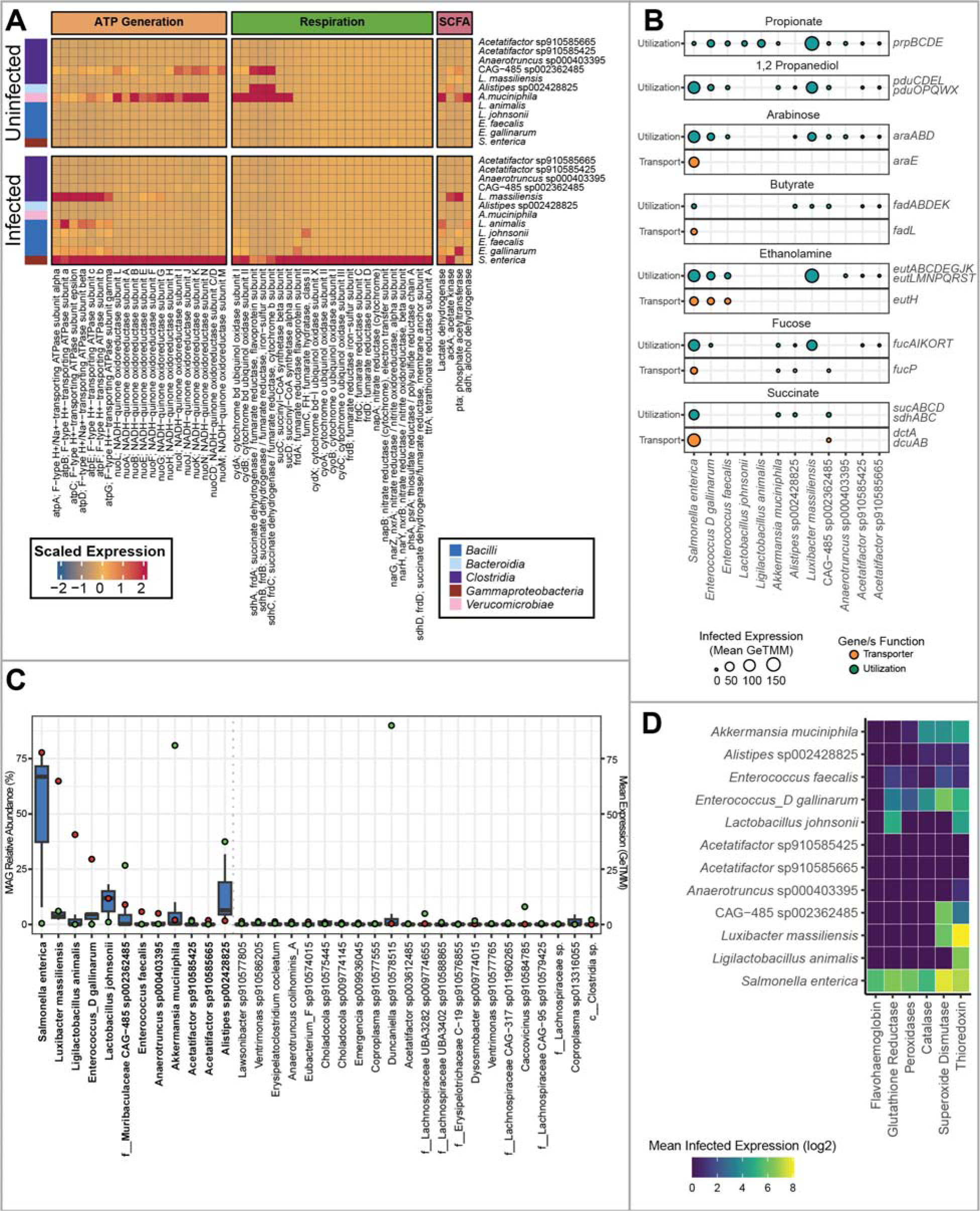
Gene expression of the most active bacteria during infection reveal substrate utilization overlap with *Salmonella.* **A** Heatmap of actively expressed genes during infection that are also significantly differentially expressed between treatments. Cell values are GeTMM scaled totals of all genes linked with a particular gene description (x-axis) averaged across samples within a treatment and expressed by an individual taxon (y-axis). Colored boxes on the left indicated gene class. **B** Carbon utilization and transport expressed by prominent bacteria during *Salmonella* infection. **C** Box plots (blue) showing the MAG relative abundance (GeTMM normalized mapped reads) of the 35 bacteria with the highest average expression (total annotated genes) in *Salmonella*-infected mice. Red points indicate average MAG expression in the infected treatment and green points indicate average MAG expression in the uninfected treatment. Taxa are ranked left to right by most detected expression during infection. Bold taxa are those with ≥ 1.5 mean expression (GeTMM) in the infected treatment, a threshold marked by the dotted gray line. **D** Average expression of oxidative stress response genes by prominent taxa during infection.

*Salmonella* respiratory processes were active at day 11, when we found evidence for aerobic respiration via expression of high and low affinity oxidases (*cydAB*, *cyoABC*, Fig. 2A). *Salmonella* also expressed genes for sulfur respiration, including the genes for tetrathionate (*ttrA*, *ttrB*, and *ttrC)* and thiosulfate reduction (*phsA*, *phsB*, and *phsC*). It is thought that these oxidized inorganic sulfur pools are created by reactive oxygen species (ROS) oxidization of hydrogen sulfide produced by host colonocytes during inflammation, a process instigated by *Salmonella* (34,35). Prior reports describe *Salmonella* with the capacity to utilize nitrate, nitrite, and trimethylamine oxidase (TMAO) (36–38) and these metabolisms (*napAB*, *narGHZY*, *nxrAB, torAC*) were also expressed in our data (Fig. 2A, Data 2). We note genes (*torA, torC, torR*) in the gene complex for TMAO reduction had detectable expression, but much lower than genes to use oxygen, sulfur, or inorganic nitrogen compounds (Data 2). To conclude, our expression data suggest *Salmonella* has the capacity to respire oxygen, sulfur, nitrogen. Our findings support the notion these myriad of respiratory strategies could provide an energetic advantage to outcompete commensal fermentative bacteria during infection (11,33,36,39,40), which had much lower expression of respiratory metabolisms (Fig. 2A). The relatively low *Salmonella* relative abundance (and presumably low inflammation) in mouse I7 at the time of metatranscript sampling (Fig. 1B) could offer one explanation for the multiple *Salmonella* metabolic strategies simultaneously detected in our expression data.

Next, we wanted to explore *Salmonella* carbon utilization as potential substrates for competition with commensal gut membership. It has been proposed that this respiratory capacity allows *Salmonella* to use lower energy compounds unavailable to many obligate fermenters, such as propionate, succinate, 1,2 propanediol, and ethanolamine as well as utilize sugars like fucose and arabinose (10,14,17,41–46). During day 11 of infection *Salmonella* expressed genes for the utilization of all the listed carbon sources, and the operons for arabinose (*ara*), 1,2 propanediol (*pdu)*, and ethanolamine (*eut*) were the most highly expressed of the carbon utilization genes examined (Fig. 2B, Data 2). We observed considerable overlap in carbon use gene expression by some commensal bacteria in infected microbiomes, indicating niche overlap with *Salmonella* that could be tunable for enhanced metabolic competition in future probiotic interventions (Fig. 2B, S2).

### Metatranscriptomics reveals diverse microbiota members are metabolically active during Salmonella infection

We were next interested in ways other present and metabolically active microorganisms persisted during *Salmonella* infection. *Salmonella* was the most relatively abundant bacteria in infected samples (50.3% average genome relative abundance) followed by *Lactobacillus johnsonii* (15% average relative abundance) (Fig. 2C, Data 3). While *Salmonella* also displayed the highest metatranscript recruitment during infection (77.7 mean GeTMM), *Luxibacter massiliensis* (*Clostridia*)*, Ligilactobacillus (*formerly *Lactobacillus B*) *animalis* (*Bacilli*), and *Enterococcus D gallinarum* (*Bacilli*), and *Lactobacillus johnsonii* (*Bacilli*) were the next most expressed lineages in infected communities (Fig. 2C). These strains are all *Bacilli*, reflecting class enrichment reported in our 16S rRNA analyses (Fig. 1B). Highlighting the discrepancy between genome abundance and transcript recruitment, *Alistipes* sp002428825 (*Bacteroidia*) and *Akkermansia muciniphila* (*Verrucomicrobiae*) were some of most dominant genomes of the infected gut (third and fourth highest proportion of the community), yet their average expression during infection was only 1.67 and 2.03 mean GeTMM respectively. We consider it possible these strains were persisting but not metabolically active at the time of our infection sampling. These findings emphasize the importance of expression in addition to genomic potential when illuminating microbial targets to successional resiliency following infection.

We found evidence that 56 MAGs other than *Salmonella* expressed metabolic genes during infection (Data 2), a phylogenetically diverse group of genomes belonging to phyla *Actinobacteria* (n=1), *Verrucomicrobiota* (n=1), *Bacteroidota* (n=4), *Firmicutes* (n=5), and *Firmicutes A* (n=45). The most active (≥ 1.5 mean GeTMM) members in feces from the *Salmonella* infected gut included 11 members of the *Lactobacillaceae*, *Lachnospiracea*, *Enterococcaceae*, *Muribaculaceae*, *Ruminococcaceae*, *Rikenellaceae*, and *Akkermansiaceae* (Fig. 2C). Consistent with our previous report, we showed interesting complexity in the *Clostridia* with some genomes persisting during infection while other dominant genera became reduced during infection (31). We note four of the top eleven most active lineages in infected mice from this study are of the *Clostridia* class (Fig. 2C) and 45 *Clostridia* MAGs expressed metabolic genes in infected metatranscriptomes (Data 2). Consistent with previous publications (15,47), the expression data here indicated butyrate production by *Clostridia* via *ptb* and *buk* was reduced during infection. Butyrate was instead provided primarily by *Alistipes* (*Bacteroidia*) and *Enterococcus faecalis* and two low abundant *Clostridia* (genera *COE1* and *Emergencia*) in the infected community. In total 52 bacterial lineages, including the most active bacteria, co-expressed some of the same genes for carbon and energy utilization as *Salmonella*, indicating community-wide metabolic overlap and exchange were occurring at this stage of infection (Figs. 2A, S2, S3, S4, Data 2). Interestingly, of the top eleven most active MAGs, eight were more active in infected mice than uninfected mice, showing how *Salmonella* infection enriches for new metabolically active community members.

### Adapting to inflammation requires mechanisms for dealing with inflammatory oxidation

To determine how enriched genomes retooled their metabolism from the non-inflamed to the inflamed infection conditions we performed a differential expression analysis at the MAG level across the two treatments. Consistent with the need to be able to respire or withstand oxidative conditions created during inflammation (Fig. 2A, S4), *Akkermansia muciniphila* and *Muribaculaceae* CAG-485 sp002362485 transcribed genes for respiring oxygen under microaerophilic or lower oxygen conditions (*cydAB*), albeit with less transcripts recruited to these MAGs in the infected than the control. Perhaps this is because of competition for molecular oxygen with *Salmonella* which expressed both low (*cyoABC*) and high affinity oxidases (*cydABX*) (Fig. 2A). We note *Akkermansia* has been reported to use NADH dehydrogenase in concert with cytochrome bd ubiquinol oxidase to respire in microaerophilic conditions (48) and both genes are expressed in our data from the infection communities (Fig. 2A, S4, Data 2). While the literature is scant concerning *Muribaculaceae*, our data suggest at least some oxygen respiration may be possible via expression of NADH dehydrogenase and high affinity oxidases by this taxon (Fig. 2A, Data 2).

Given the breadth of respiratory metabolism expressed by *Salmonella*, we were interested if other dominant and persisting members utilized similar energetic strategies. Surprisingly, we failed to detect expression of denitrification or sulfur respiration pathways in the top infection-active taxa. Yet, gene expression profiles of *Enterococcus D gallinarum* and *Lactobacillus johnsonii* indicated fumarate reduction increased during infection (Fig. 2A, S3, S4). While some *Enterococci* have the capacity for aerobic respiration, our genome lacked an electron transport chain, terminal oxidase, and complete tricarboxylic acid cycle despite being 96% complete, suggesting that respiration of oxygen and or fumarate is unlikely. Alternatively, strains of *Enterococcus* can use fumarate under anoxia to enhance growth during glucose fermentation, likely due to its role as a hydrogen sink (49). Further physiological studies are needed to support this supposition; however, the metatranscript data suggest fumarate may be an important compound establishing *Enterococcus* and *Lactobacillus* enrichment in the infected microbiome, potentially providing a metabolic advantage over strictly fermentative commensal membership. Along these lines of unexpected growth advantage during infection, the two most transcriptionally active bacteria in infected mice (*Luxibacter massiliensis, Ligilactobacillus animalis*) showed significantly higher expression of genes for F-type proton pumps (Fig. 2A), indicating ATP generation via proton motive force as a strategy during inflammation.

Further, we considered it likely that many persistent commensal bacteria employed mechanisms to tolerate inflammatory conditions during infection. The most transcriptionally active genomes in the infected community all expressed multiple oxidative stress response genes (Fig. 2D). Of these various strategies, the most common across the community was thioredoxin, where thioredoxin-dependent peroxiredoxin (*tpx*) reduced reactive oxygen species (ROS) in a manner dependent on thioredoxin 1 (*trxA*) (50). Catalases, peroxidases, and superoxide dismutase were also prevalently expressed amongst the most active taxa, as well as glutathione reductase (*gor*) (Fig. 2D). *Gor* is important to LAB for maintaining reduced glutathione amidst oxidant stress from host immunity (51). In summary, the ability to respire or withstand oxidative stress seems an important feature expressed by transcriptionally active members of the microbiome during infection.

### Intertwined lactate metabolisms are expressed during Salmonella infection

Our expression analysis indicated lactate was a possible energy source used by *Salmonella* (Figs. 2A, S3). Here we show this metabolite was likely important community-wide, with eleven of the most transcriptionally active genomes in the infected intestine expressing lactate dehydrogenase genes (Fig. 3). While studies using *Salmonella* genetics in gnotobiotic or reduced complexity microbiomes reported that L-lactate from colonocytes was the primary lactate enantiomer used by *Salmonella* (9,52), our metatranscriptome data offered an opportunity to track the production and consumption of D- and L-lactate by the microbiome as lactate dehydrogenases can be specific for different lactate enantiomers – *ldh* (L-lactate dehydrogenase, K00016), *lldD* (L-lactate dehydrogenase, K00101), *dld* (D-lactate dehydrogenase, K03777), and *ldhA* (D-lactate dehydrogenase, K03778). These gene products catalyze either the production or the consumption of lactate to form other metabolic end products. The directionality of these reactions is inferred by gene type, energy favorability, and metabolic capabilities of closely related characterized strains (9,52,53). We exercised caution determining sources and sinks of lactate from gene expression as many microorganisms like LAB (e.g. *Lactobacillus* and *Enterococcus* genera) produce lactate when fermenting, while bacteria like *Clostridia* can ferment lactate to butyrate or acetate during fermentation, and *Salmonella* can oxidize lactate when respiring (53). As such, a combination of genomic context and known characterizations of closely related physiological isolates was used to deduce possible lactate production and consumption in the *Salmonella*-included microbiome.

**Fig. 3:**
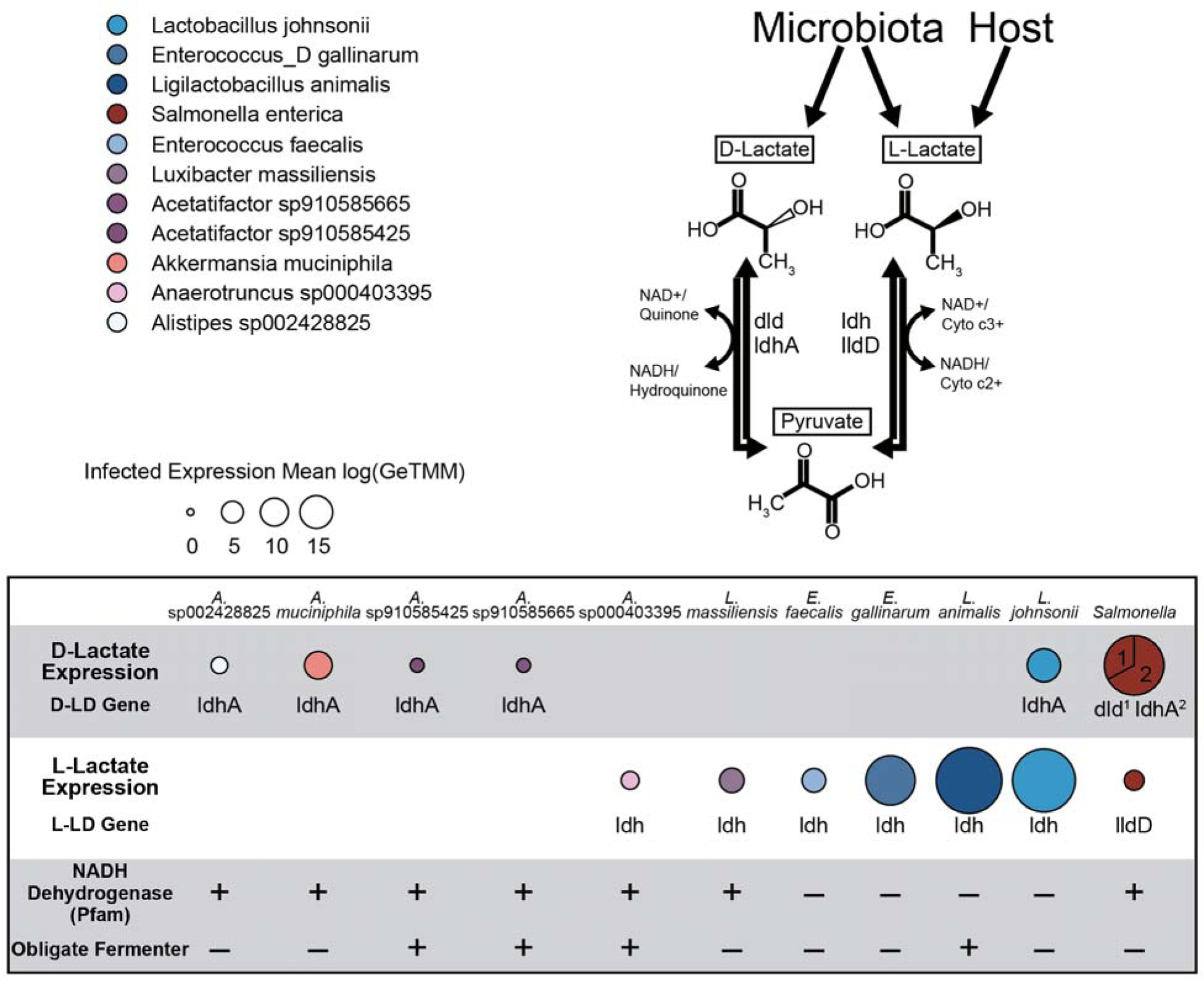
*Salmonella* lactate dehydrogenase expression reveals both D-lactate production and oxidation by important membership in the inflamed gut. Expression of individual lactate dehydrogenase genes by select taxa during *Salmonella* infection. Circle sizes indicate average log expression in infected samples recruited by individual genomes or averaged within higher taxonomic groups. Numbered sections in the *Salmonella* pie chart denote the proportion of total lactate dehydrogenase expression represented by each gene. NADH dehydrogenase gene content was examined for each MAG, a plus (+) indicates at least one Pfam annotated gene is present, a minus (-) indicates no genes were annotated “NADH dehydrogenase”. Positive (+) obligate fermenter designation was assigned if a MAG showed no expression of any respiratory genes as surveyed in Fig. 2. Negative (-) obligate fermenter designation was applied to all other MAGs in the figure and is supported by literature discussed in the text.

Here we report *Salmonella* expression of genes for D-lactate exceeds that for L-lactate (Fig. 3). *Salmonella* was also the only organism expressing both lactate dehydrogenases for D-lactate (*dld*, *ldhA*, Fig. 3), and given its respiratory capacity and its known use of lactate as a substrate (54), we infer D-lactate consumption by *Salmonella*. Though many factors may influence gene expression, these findings show D-lactate likely produced by the microbiota, as well as L-lactate as previously thought to be colonocyte derived are both possible substrates used by *Salmonella* within an intact microbiome. We also detected the lactate metabolite in 3.9-fold greater mean abundance in the infected mice than in the non-infected mice (Supplementary Data 3), however our mass spectrometric methods lacked enantiomeric resolution.

We next examined if microbial community expression supported a model where D-lactate was produced by members of the microbiota. Of the most prominent infection community members, *Lactobacillus johnsonii, Alistipes* sp002428825, *Akkermansia muciniphila*, and two *Clostridia* had detectable levels of D-lactate dehydrogenase (*ldhA*) expression in the infected samples (Fig. 3). Outside of *Salmonella*, the highest *ldhA* expression was from *Lactobacillus johnsonii*, which displayed a significant 3.48 log fold-change increase of *ldhA* expression during infection, likely due to its nearly 38-fold enrichment under the same conditions (Fig. 3, S4, Data 1-3). Of the D-lactate metabolizing strains, we confirmed *Lactobacillus johnsonii* lacked an electron transport chain for respiration suggesting lactate was a metabolic product of fermentation (55), and conversely, the other strains all have NADH dehydrogenase genes (Fig. 3). Furthermore, physiological characterization of isolates closely related to this MAG demonstrated that *Lactobacillus johnsonii* produces both D and L-lactate (56), often times in a 60/40 (D/L) racemic proportion (56,57), further supporting the notion that lactate was likely produced by this microorganism (58). Prior work deduced *Salmonella* preference for host derived L-lactate in gnotobiotic mice (9), and we posit *Salmonella* alters its gene expression for lactate utilization based on competition with other members of the microbial community and greater D/L enantiomer proportion from LAB activity in the infected microbiome.

Most of the L-lactate dehydrogenase gene expression occurred in *Bacilli*, especially LAB like *Lactobacillus johnsonii* and *Ligilactobacillus animalis*, which are known to produce L-lactate (53,56,59). We show this production is important during infection, where L-lactate dehydrogenase (K00016) was upregulated 1.8 and 3.9 (mean log fold-change) respectively in the infected treatment (Fig. 3, S3, Data 2). *Enterococcus* genera *gallinarum* and *faecalis* also highly upregulated K00016 during infection 4.3 and 3.8 mean log fold-change (Fig. 3, S3, Data 2). Taken together we consider it likely that in addition to colonocytes, the microbiota, especially LAB, are another source of L-lactate during infection. Alternatively, while they may be producing L-lactate, we cannot rule out that other fermentative members active during infection, like members of *Clostridia* (e.g., *Luxibacter, Acetatifactor* spp., and *Anaerotruncus*) may instead ferment lactate, as demonstrated by other members of this class (53). These results, along with the *Salmonella* data, point to lactate cross feeding as potentially important for persistence during inflammation.

Combined our expression and genomic data expands the current understanding of lactate metabolism in the infected microbiome. Our metabolite and metatranscript data indicate lactate, both host-derived and microbiota-derived (specifically metabolically active LAB), is an important metabolic currency in the infected gut. It is likely that *Salmonella* utilizes both lactate enantiomers and that strains of *Akkermansia*, *Alistipes*, and *Clostridia* may also compete with *Salmonella* for lactate in the inflamed gut. We note that fermentation of lactate by *Clostridia* would result in butyrate and acetate production, which would be important in facilitating short chain fatty acid availability for the host after clearing an infection.

### Metabolites from the infected microbiome indicate the importance of organic sulfur pools in the gut

To further expound relevant processes in the *Salmonella*-infected microbiome we performed untargeted metabolomics and compared the metabolomes in uninfected mice and those at the later stages of infection. Broadly, we noticed a significant (PERMANOVA p=0.001) difference in chemical composition of feces from mice 12 days post infection and feces from control mice collected at the same timepoint (Fig. 4A, Data 4). Principal component analysis revealed strong clustering of metabolomes by treatment due largely to differences in abundance of flavonoids, amino acids, and bile acids (Fig. 4A). We found that many sulfonated compounds including sulfated flavonoids, bile acids, and sulfur containing amino acids like taurine and methionine differed in abundance between treatments (Figs. 4A, B, Data 4). Sulfonated bile acids were significantly (ANOVA, p<0.05) more abundant in infected guts, as were oxidized methionine and sulfonated flavonoid species, while uninfected mice harbored higher abundances of deconjugated bile acids and free taurine (Fig. 4C).

**Fig 4:**
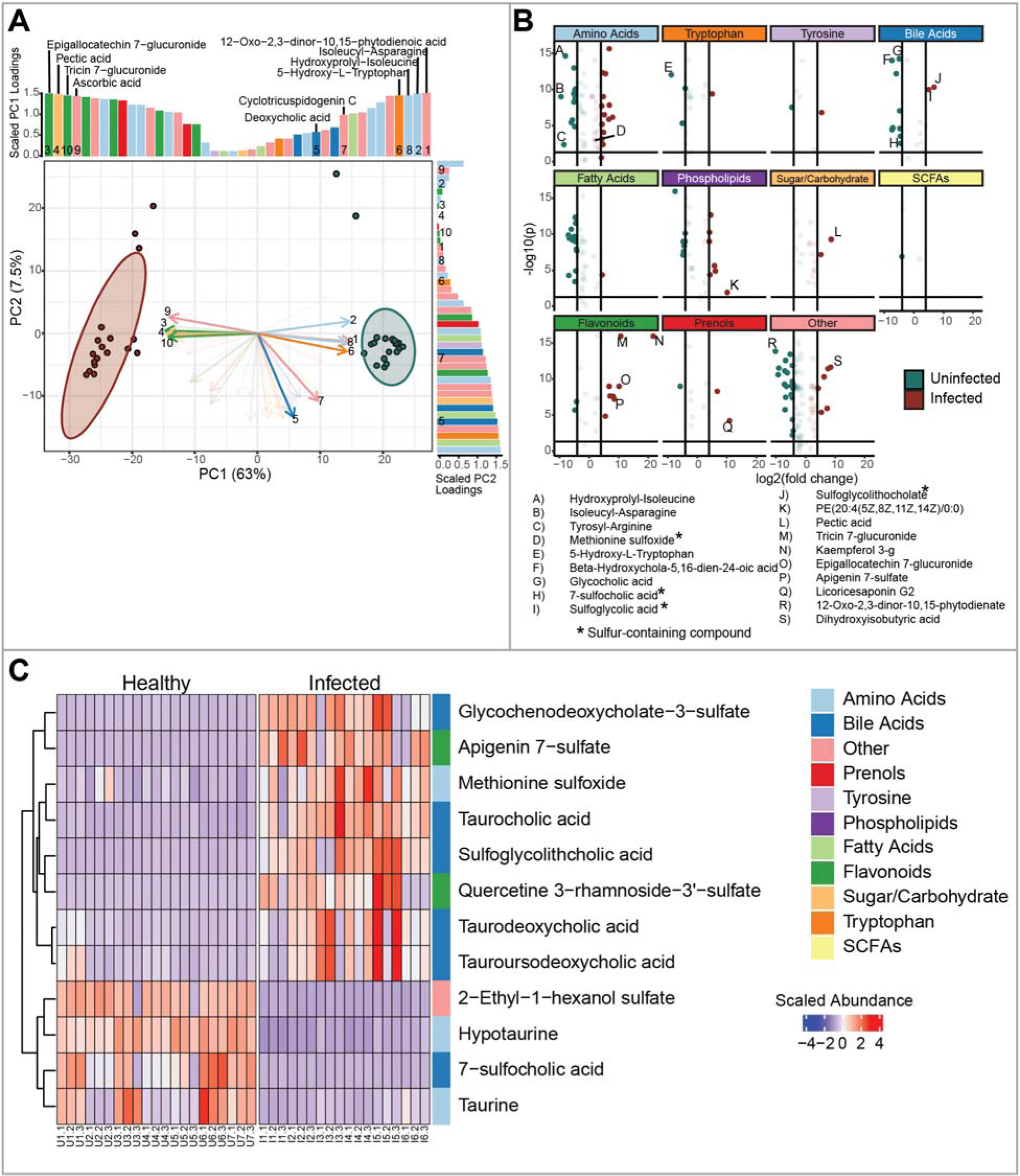
Uninfected metabolomes differ from inflamed microbiomes where sulfonated bile acids and flavonoids are prevalent. **A** Principal component analysis (PCA) of metabolites from infected and uninfected mice colored by treatment (red = infected, green = uninfected). Arrows and bars are significant (ANOVA p < 0.05) compounds. Dark arrows are the top 10 compounds that explain the PCA ordination variance (Euclidian distance from tip to centroid). Bars are arranged by absolute value of compound loading for each component, and they are colored by compound group. **B** Volcano plots of significant metabolites separated by compound group. Points are colored by treatment (red = infected, green = uninfected) and dark points indicate significant (ANOVA p < 0.05) compounds with at least 4x log2 fold change between treatments. Select compounds (most changed or system relevant) are labeled below the plot and sulfur containing compounds are denoted with an asterisk. **C** Heatmap showing the center scaled abundance of significantly different sulfur containing compounds in each treatment.

The organic sulfur pool consisting of sulfated bile acids, sulfated flavonoids, and methionine sulfoxide in infected mice, provides a potential source of inorganic sulfur via commensal and *Salmonella* activity (Figs. 4C, 5). Flavonoids and bile acids are conjugated to sulfonate in the liver and released back to the gastrointestinal tract via enterohepatic circulation, acting as host derived sources of sulfur in the inflamed gut (60,61). One of the most discriminant sulfur compounds in our metabolome was methionine sulfoxide which is the oxidized form of methionine and may originate from protein in the diet or dead microbial cells (62). One report describes the importance of methionine to *Salmonella* virulence, remarking how significant it is that *Salmonella* encodes redundant mechanisms for its acquisition or production (63), and so we chose to examine pathways for utilization of this amino acid more closely in our data.

**Fig 5:**
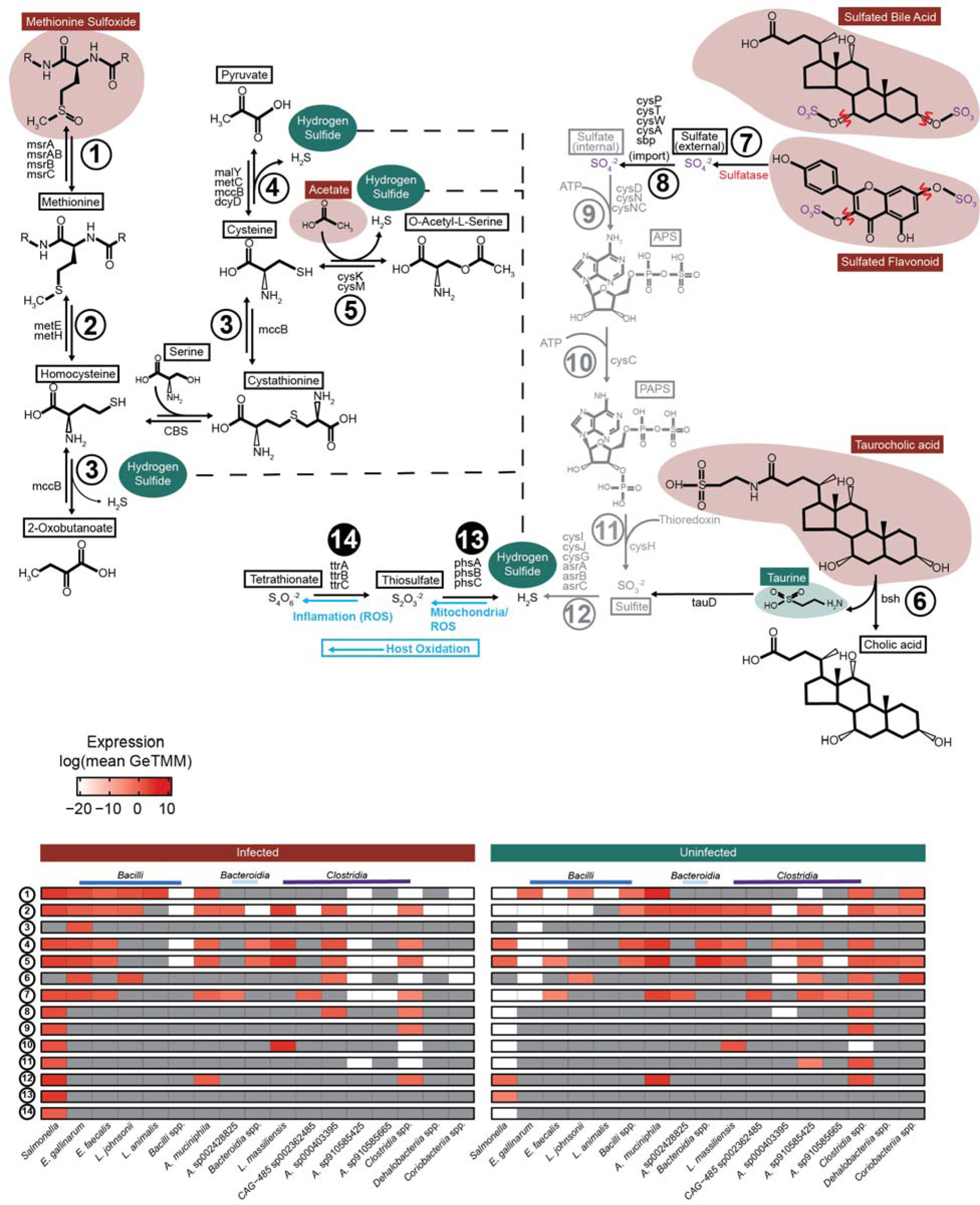
Infected community expression implicates commensal membership with hydrogen sulfide production and *Salmonella* sulfur metabolism. Pathways depicting amino acid (left) and sulfated bile/flavonoid compounds (right) as organic sulfur substrates used by the infected microbial community to produce hydrogen sulfide. Numbered pathway arrows correspond with numbered rows in the heatmaps. Dashed lines connect sections of the amino acid pathway to the hydrogen sulfide pool. Blue arrows show host oxidation of inorganic sulfur. Sulfate (purple) released by sulfatase activity (indicated in red) precedes sulfate reduction steps following ABC transporter-mediated translocation colored gray. Compounds outlined in red and green are significantly more abundant in infected metabolomes or uninfected metabolomes respectively, and compounds outlined in black are not present in our metabolite data. The heatmap on the right indicates pathway expression in uninfected mice and the heatmap on the left indicates expression from infected mice. Cells are colored by average log expression and are dark grey if no expression was detected. Pathway steps denoted with black filled circles are specific to *Salmonella* in the community.

### Oxidized inorganic sulfur made available by the gut microbiota exclusively supports *Salmonella* sulfur respiration

Our metabolite data revealed that methionine sulfoxide was a key discriminant metabolite, more prevalent in infected microbiomes with *Salmonella*. We hypothesized that this organic sulfur source, and others derived from it, could serve as a source for sulfur exchanges that ultimately lead to *Salmonella* respiration. Given that ROS can oxidize hydrogen sulfide yielding *Salmonella* respiratory compounds tetrathionate and thiosulfate, we were most interested in the steps to reduced sulfur from methionine sulfoxide. Outlined in Figure 5 is the initial conversion of methionine sulfoxide to methionine (step 1), then to homocysteine (step 2), ultimately yielding 2-oxobutanoate, pyruvate, or O-acetyl-L-serine and hydrogen sulfide (steps 3, 4, and 5). Many active members of the inflamed gut expressed the first two steps in this pathway – step 1 methionine sulfoxide reductase (*msrABC*) and step 2 homocysteine methyltransferases (*metEH*). *Salmonella*, *Akkermansia*, and most predominant *Bacilli* actively expressed these genes, showing methionine sulfoxide is likely an important sulfur source in the inflamed gut. However, only the *Enterococcus D gallinarum* genome encoded and expressed genes for cystathionine-γ-lyase (*mccB*) to convert homocysteine to 2-oxobutanoate and cysteine to pyruvate, both reactions that release hydrogen sulfide and both only expressed in the infected gut (64).

Genes involved in cysteine metabolism (step 4: *dcyD*, *malY*, *metC*, and *mccB*), as well as cysteine synthase (step 5: *cysK*, *cysM*) were examined, as both these steps could release hydrogen sulfide from cysteine degradation (64). These pathways were more prevalently expressed than the complete methionine pathway (Fig. 5). *Salmonella*, *Akkermansia*, *Enterococcus*, and *Clostridia* members including *Luxibacter* and *Anaerotruncus* expressed both steps 4 and 5 at high levels to release hydrogen sulfide during infection (Fig. 5). In summary, our paired expression and metabolite data provide compelling evidence that sulfur-containing amino acids like cysteine and methionine may be an understudied source of microbially derived hydrogen sulfide, suggesting a larger role of commensal membership in sulfur cycling in the inflamed microbiome.

Additional organic sulfur prevalent in our infection data and also discriminate between treatments were sulfated bile acids (e.g. Sulfoglycolithocholate, Glycochenodeoxycholate-3-sulfate) and flavonoids (e.g. Quercetin 3-rhamnoside-3’-sulfate, Apigenin 7-sulfate). It is recognized that *Salmonella* and other gut microbes express sulfatases that desulfurize these compounds releasing sulfate (Fig 5, Data 2) (65). Sulfatase expression by the twelve most active bacteria (bar *Akkermansia*) was 2-fold greater in the inflamed gut, and *Enterococcus faecalis, Enterococcus D gallinarum,* and *Salmonella* accounted for 94% of all sulfatase expression in the infected microbiome (Fig. 5). *Akkermansia* sulfatase expression dramatically decreased in the infection treatment (45.8-fold) (Fig. 5). Overall, these data contribute to a growing body of work aimed to better understand bacterial production of sulfatase in the mammalian colon and its implication on microbiome structure and nutrient acquisition from host-derived and diet-derived sources (66,67). Our data places emphasis on *Enterococcus* activity transforming the gut organic sulfur pool generating a source of free sulfate during *Salmonella* infection.

Reduction of sulfate (generated from sulfatases) could also lead to production of hydrogen sulfide. Sulfate reducing bacteria have been reported to reduce up to 18.6 mmol of sulfate per day in the mammalian gut from organic sources like bile acids and mucins (64). Sulfur is reduced in the gut either via the dissimilatory sulfur reduction (DSR) pathway (*sat, aprAB, dsrAB*) or the assimilatory sulfur reduction (ASR) pathway (*cysDNCHIJ, asrABC*) (64,68–70). We detected no DSR gene content in the CBAJ-DB v1.2. Instead, we found genes for ASR expressed by both *Akkermansia* and *Salmonella* (step 12, Fig. 5), but only *Salmonella* contained genes in the four other steps for importing and converting extracellular sulfate to sulfite, all of which were highly expressed during infection (Fig. 5). Additionally, three *Clostridia* lineages expressed either *asrA, asrB,* or *asrC* to catalyze the same reduction (Fig. 5) (64,69). Our gene data suggest ASR enzymes may be important contributors to sulfate and sulfite reduction, more so than canonically studied DSR processes. These findings from the infected mouse gut are supported by a study in human gut microbiota, where ASR genes were twice as abundant as DSR genes and were deemed critical modulators in colon cancer (64).

Here we have shown two distinct mechanisms for hydrogen sulfide generation in the gut using microbiota driven 1) amino acid degradation and via 2) removal of sulfate from sulfated organic compounds like bile salts and subsequent reduction via anaerobic sulfite reductase. Host catalyzed sulfur oxidation would generate tetrathionate and thiosulfate in the infected gut, as previously reported (11,20,34), compounds which our data show *Salmonella* is uniquely positioned to respire (steps 13, 14 in Fig. 5). Mining of CBAJ-DB v1.2 genomes for these sulfur respiratory capacities revealed only *Salmonella* had capacity for both tetrathionate and thiosulfate reduction. Two other *Coriobacteriia* lineages had genes to reduce thiosulfate, yet combined, these lineages comprised less than 0.001% of the infected community (Data 3). One other lineage (*Adlercreutzia muris*) besides *Salmonella* expressed thiosulfate reduction genes during infection, yet at a rate 50-fold lower than *Salmonella*. Likewise, both tetrathionate and thiosulfate metabolisms were not expressed in the uninfected gut (Data 2), confirming the metabolic specialization of this pathogen for microbiota, diet, and host derived metabolic products.

Our findings underscore the complex interplay between pathogen, diet, microbiota, and host in the murine gut model. Our results are supported by a growing body of research in humans, recognizing the importance of a balanced sulfur cycle to gut health, where imbalances have been linked to inflammatory bowel disease and colorectal cancer (64,71). It is likely that the sulfur metabolisms shown here by metabolite and metatranscriptomic analysis could possibly be more amplified in the westernized human diet compared to the mouse. For example, western diet studies using fecal homogenates demonstrate higher production of hydrogen sulfide from organic sulfur amino acids compared to inorganic sources, and high protein intake increased the sulfur-free amino acid content of the gut (72,73). Future work is needed to evaluate the diet-microbiota-pathogen linkages in more realistic diet regimes while also extending and curating sulfur metabolism more holistically in the human gut microbiome.

### Conclusions

The work presented here has identified a set of commensal bacteria that persist during *Salmonella* inflammation and infection to actively participate in the utilization and production of key metabolites in the inflamed gut ecosystem. These results offer multiple exciting avenues for probiotic strain identification of bacteria robust to enteric inflammation and pathogen perturbation that may be pivotal to reestablishing normal gut function following infection. We confirmed *Salmonella* expression of genes to utilize lactate, tetrathionate, and thiosulfate *in vivo* amidst an intact microbial community. Our data also expands current paradigms of *Salmonella* lactate utilization during infection and proposes a role for commensal bacteria in *Salmonella* sulfur and lactate use. Additional work is necessary to confirm the mechanistic assertions made here, but the inferences drawn underpin the value in holistic examination of the microbiome afforded by a multi-omics approach.

## Methods

### Strains and media

*Salmonella enterica* serovar Typhimurium strain 14028 (*S. typhimurium* 14028) cultures were washed and resuspended in water after overnight incubation in Luria-Bertani broth at 37 °C with constant agitation.

### Animals and experimental design

Female CBA/J mice were procured from The Jackson Laboratory (Bar Harbor, ME). Mice were randomly selected to populate cages and treatment groups (infected n=44, uninfected n=23), and were kept 5 per cage in conventional enclosures in a temperature controlled 12-hour light/dark cycle. Irradiated mouse chow (Teklad, 7912) was made available *ad libitum* to all mice. Mice in the infected group were inoculated with 10^9^ CFU *S. typhimurium* 14028 via oral gavage on day 0 with no subsequent treatment. Uninfected mice were left with no treatment. Animal experiment protocol was approved by The Ohio State University Institutional Animal Care and Use Committee (IACUC; OSU 2009A0035).

Mouse fecal pellets were collected from thirty-eight mice before and after treatment initiation (on days −2, −1, 0, 10, 11, or 12) on autoclaved aluminum foil. Sample details by mouse are outlined in Supplementary Data 3 and Figure 1. Fecal pellets were immediately placed in labeled microcentrifuge tubes and flash frozen with EtOH/dry-ice prior to storage at −80 °C until further processing.

### DNA/RNA Extraction and Sequencing

All nucleic acid extraction for 16S rRNA amplicon sequencing was performed using ZymoBIOMICS DNA/RNA Miniprep Kit (Zymo Research) and stored at −20 °C until further processed. PCR amplification of the V4 hypervariable region of 16S rRNA gene was performed using 30 cycles and unique sequence barcodes in each primer were used to identify multiplexed samples. Both primers (universal 515F and 806R) contained sequencer adapter regions. Total nucleic acid extraction and sequencing for metagenomics was performed as described in Leleiwi et.al. (31) at the Genomics Shared Resource facility at Ohio State University.

Metatranscriptomic RNA extraction and isolation was performed on feces from 10 mice sampled 11 days post infection with Zymo-Seq Ribo Free Total RNA Library Kit Cat No. R3000 and 2 x 151 bp paired end reads were produced from corresponding cDNA at UC Denver Sequencing Facility using an Illumina HiSeq 2500 Sequencing System.

### Metabolite Sample Preparation and Analysis

1 mL of a solution composed of three different solvents (water/methanol/dichloromethane, 1/2/3, v/v/v) was used for fecal metabolite extraction, followed by physical disruption with a sonicator (Bioruptor®, Diagenode, Belgium). The disrupted fecal suspension was vortexed and incubated at room temperature. The top aqueous layer was transferred, dried under vacuum, and reconstituted for metabolomics analysis.

The Agilent 6545 QTOF mass spectrometer equipped with Agilent 1290 UHPLC systems liquid chromatography was used for an untargeted metabolomics study on mouse fecal pellet extractions (n=13) from day 12. To improve the metabolite coverage, the prepared fecal samples were subjected to reversed-phase liquid chromatography with C18 column (ACQUITY UPLC® HSS T3 1.8 µm, 2.1 x 100 mm, Waters Corporation, MA, USA) and hydrophilic interaction liquid chromatography (HILIC, ACQUITY UPLC® BEH HILIC 1.7 µm 2.1 X 150 mm, Waters Corporation, MA, USA). Water with 0.1% formic acid (A) and acetonitrile with 0.1% formic acid (B) were used for reversed-phase separation. The flow rate was set at 0.3 mL/min with the gradient as follows: 2% B for 0-2 min, from 2% B to 30% B for 4 min, to 50% B for 8 min, and 98% B for 1.5 min and held at 98% B for 1min, then returning into initial gradient for equilibrium for 1.5 min. For HILIC separation, Water/acetonitrile (95/5) with 0.1% formic acid and 10 mM ammonium formate (A) and water/acetonitrile (5/95) with 0.1% formic acid and 10 mM ammonium formate (B) were prepared. For gradient elution, 99% B was held for 2 min, gradually reduced to 75% B for 7 min and reduced again to 45% B for 5 min. And the gradient was held at 45% B for 2 min, and returned to 99% B. The flow rate was set at 0.3 mL/min. The quality control (QC) sample was prepared by mixing an equal volume of each sample. The QC sample was analyzed after every 6 samples.

The collected mass spectra were converted into mzML format with MSconvert in Proteowizard (74). The data analysis for untargeted, global profiling experiments was performed with Progenesis QI (Waters Corporation/Nonlinear Dynamics). The Compound MS/MS library used in the study for metabolite annotation was the embedded Progenesis database with METLIN/Waters collaboration for fragmentation patterns, human metabolome database (HMDB), E. coli metabolome database (ECMDB), and Lipid Maps. All annotated metabolites were manually inspected to be reported and metabolite significance between treatments was determined with ANOVA (Data 4).

All metabolite samples were included in the ANOVA significance calculation (uninfected n=7, infected n=7) while later analyses omitted the metabolome from mouse I7 in the infected group because it was not a high-responder. Missing values in the remaining metabolite data were imputed with half the smallest value recorded for each feature. Data were log transformed and Pareto scaled (75) and principal component analysis was performed in R version 4.1.3 with the prcomp function in the vegan package (v2.6-4). Permanova was performed on Bray-Curtis distances calculated from untransformed data using the vegdist function from vegan. Euclidian distances from the centroid were calculated for important (threshold >0.09 & <-0.09) loadings in PC1 and PC2 (see script Metab_pca.R).

### 16S rRNA Gene Analysis

Amplicon sequencing fastq data generated for this experiment were a subset of data encompassing days −2, −1, 0, 10, 11, and 12 from a larger experiment under Bioproject number PRJNA348350 and were processed in a QIIME2 2019.10.0 environment, with reads demultiplexed and then denoised with DADA2 (76,77). For all sequencing runs (n=6), forward reads were truncated at 246 bps and reverse reads were truncated at 167 bps. Feature tables from each sequencing run were combined and ASVs were later assigned taxonomy with the silva-138-99-515-806-nb-classifier in a QIIME2 2022.8.0 environment (76,78). The resulting ASV table was filtered with R version 4.1.3 to include only samples from days within the scope of this paper and to remove any samples with zero counts for every ASV (n=2 samples). Next, ASVs were removed with zero counts in every remaining sample (n=22,175 ASVs), and subsequently samples with fewer than 1000 counts across all remaining ASVs were dropped (n=3 samples). Finally, any ASVs designated as mitochondria, chloroplast, unassigned at the domain level, or assigned Eukaryota were removed (n=66 ASVs). The 16S V4 region from *Salmonella enterica* Typhimurium ATCC 14028 reference genome was manually aligned with Geneious Prime® 2020.1.2 to a single *Enterobacteriaceae* ASV (ASV ID = 4cbfff144d4e7a4e0f4619ed505be070) with 100% sequence identity to confirm taxonomy as *Salmonella*. The designation “high-responder” was applied to any mouse with *Salmonella* ASV relative abundance ≥ 25% in at least one sample. Any mouse from the infected group was not included in the analysis unless it was a high-responder or had additional sampling for metatranscriptomic, metagenomic, or metabolomic analysis. The resulting feature table contained 1405 unique ASVs (Data 3).

### 16S rRNA Statistical Analysis

#### ASV Community Metrics and Class Significance

ASV communities from mice with metabolomics data and metatranscriptomics data (n=14) were categorized by sampling day: Early (day −2 – day 0) and Late (day 10 – day 12). Average ASV relative abundance within each class was calculated and the 8 most abundant classes were retained and any other ASVs were classified as “Other”. Significant differences between classes and treatments were calculated with the Mann-Whitney U test in R.

#### 16S rRNA Linear Discriminant Analysis

The filtered ASV table was limited to samples (n=41) from mice that also had metabolomics or metatranscriptomic data including only samples from days 10, 11, and 12. ASVs with ambiguous genus designations were removed and then relative abundance of each ASV was calculated within samples. The table was collapsed to the genus level and linear discriminant analysis was performed with LEfSe to determine important taxa from each treatment (79).

#### ASV High-Responder Correlation Network

First the filtered ASV feature table was collapsed to the genus level and features were removed with 5000 or fewer counts and ambiguously named genera were removed (“uncultured”, “uncultured bacterium”, “unidentified”). The genus table was further curated to include only data from high-responder mice collected from days 10, 11, and 12. Any genus with 1000 or fewer counts from the subsequent table was also removed from further analysis. Spearman correlation between genera was performed with the Hmisc package (v.4.7-1) in R version 4.1.3. Significant (p<0.05) positive interactions from the resulting correlation matrix were retained. Any genera with a positive correlation coefficient with *Salmonella* were further considered. The resulting correlation network was plotted with R base plot function.

### Metagenome Assembled Genome Database

A total of 3,667 (n=160 dereplicated quality) metagenome assembled genomes (MAGs) were used in this study. These MAGs were 1) obtained from the CBAJ-DB (n=2,281) with methods described in Leleiwi et.al. (31), and 2) were reconstructed from metagenomic sequencing of 6 additional mice with methods described here. This publication is the first description of MAGs originating from high fat diet metagenomes.

All metagenomic reads were checked for quality with FastQC (v0.11.9) and trimmed of low quality reads and adapters, and mouse reads were removed using BBDuk (ktrim=r, k=23, mink=11, hdist=1, qtrim=rl, trimq=20, minlen=75, maq=10) from BBTools (v38.89, https://jgi.doe.gov/data-and-tools/bbtools). Metagenomic samples (n=6) were assembled independently with Megahit (v1.1.1) and the resulting assemblies were filtered to include only contigs ≥2500 bps and binned with Metabat2 (v2.12.1,-verysensitive). Collectively, all assemblies resulted in 1,386 MAGs, which were dereplicated with CBAJ-DB using dRep (v2.6.2,-sa 0.99-comp 50-con 10), resulting in 160 dereplicated MAGs which are the basis for this paper. The dereplicated MAG set (n=160, CBAJ-DB v2.1) was used for mapping metagenomic and metatranscriptomic reads. MAG taxonomy was assigned using GTDB-Tk (v.2.1.1, r207). MAG quality was confirmed with CheckM (v.1.1.2). MAGs were mined for 16S rRNA sequences and matched to ASVs using The Microbial Ecosystem Lab software (https://github.com/WrightonLabCSU/join_asvbins). Briefly, join_asvbins workflow searches MAGs for candidate 16S rRNA sequences using MMseqs2 and a reference database (silva-138-99-seqs-515-806) and it uses Barrnap (v0.9) to predict 16S rRNA sequences. The candidate sequences are then compared to amplicon sequenced ASV sequences using MMSeqs2.

Gene annotations for each MAG in the database were produced with DRAM (v1.4.0) (80). Sulfur metabolism was curated from DRAM annotations, with manual curation of cysteine and serine sulfatase activation signatures - (C/S)*XPX*R and CXAXR in R version 4.1.3 (81). The resulting gene set was further curated to include only genes with a sulfatase motif and Pfam Sulfatase annotation, either sulfatase PF00884.26, or the sulfatase modifying factor enzyme PF03781.19.

### Metagenomic Mapping

Raw paired-end metagenomic sequencing reads from *Salmonella*-infected mice used to create the CBAJ-DB v2.1 database and one additional *Salmonella*-infected metagenome (n=7) were mapped to the CBAJ-DB v2.1 database using Bowtie2 (v.2.4.5)(82). Prior to mapping, raw reads were quality trimmed with Sickle (v.1.33) to remove sequences with phred quality scores below 20. Additionally, reads were filtered with RQCFilter2 to remove adapter sequences, RNA artifacts, phiX kmers, rRNA sequences, and any sequences that map to mouse, cat, dog, human, or common microbial contaminate genomes. BAM files produced during mapping were sorted by sequence name and filtered to include only mappings with 95% identity with Samtools (v.1.9) and reformat.sh (https://github.com/BioInfoTools/BBMap/blob/master/sh/reformat.sh) respectively. Gene counts were produced for each MAG in the database with Featurecounts (v.1.5.3) and the resulting counts table was GeTMM normalized (83).

### Metatranscriptomic Sequencing Analysis and Mapping

Raw metatranscriptomic paired fastq data collected on day 11 from five CBA/J mice in each treatment were quality trimmed to remove reads shorter than 75 bps and of lower quality than phred 20 with bbduk.sh (https://github.com/BioInfoTools/BBMap/blob/master/sh/bbduk.sh). PhiX and rRNA sequences were removed with bbmap.sh (https://github.com/BioInfoTools/BBMap/blob/master/sh/bbmap.sh) at minid=0.90. The resulting reads were then mapped to the concatenated quality non-redundant MAG database (CBAJ-DB v1.2) using Bowtie2 (v.2.4.5) using flags -D 10 -R 2 -N 0 -L 22 -i S,0,2.50. Binary alignment files were filtered to only include mappings with high sequence identity (≥97%) using reformat.sh (https://github.com/BioInfoTools/BBMap/blob/master/sh/reformat.sh) and were sorted by sequence name with Samtools (v.1.9). Individual gene counts were produced with Featurecounts (v.1.5.3) and the resulting counts table was GeTMM normalized (83). The count table was further filtered to remove any genes with 0 counts in every sample. Rarification to 2.5 Gbps was performed with reformat.sh from the BBTools suite on each metatranscriptome and the mapping process was repeated. Active MAG comparison between rarified and unrarified mapping was determined by considering genomes with at least one count in 3 or more samples. Differential gene expression and significance of GeTMM normalized filtered transcript counts was produced with limma (v.3.50.3) and edgeR (v.3.36.0) R packages. Limma analysis was performed on all metatranscriptomic samples bar that from mouse I7 which was omitted from the infected group because it was not a high-responder. The DRAM distillate and product outputs were used to group counts assigned to individual MAGs into functional categories and to determine MAG substrate utilization and individual gene expression in MAGs of interest. Metabolic gene expression was determined by a total count value greater than 0 of genes with either a Kegg ID or CAZY ID that were also classified in the DRAM metabolism summary sheets: “carbon utilization”, “Energy”, “Organic Nitrogen”, and carbon utilization (Woodcroft).

## Acknowledgements

Authorship would like to acknowledge Tyson Claffey and Richard Wolfe as integral members of the Colorado State University server management group that made this work possible; Sandy Shew is acknowledged here as primary management for the computing resources retained from The Ohio State University Unity cluster; Finally, from the Genomics Shared Resource Core at The Ohio State University Comprehensive Cancer Center for management of metagenomic sequencing, Dr. Pearly Yan is acknowledged.

## Data availability

Sequencing data supporting the results shown here are provided in the National center of biotechnology Information (NCBI) under Bioproject number PRJNA348350. The amplicon sequencing ASV table is provided in the supplemental data and CBAJ-DB v1.2 MAG fastas are provided on Zenodo https://doi.org/10.5281/zenodo.8395759. Scripts used in data analysis can be found at https://github.com/ileleiwi/Salmonella-Multiomics-Paper.

## Author Contributions

IL, KK, YK, MB, MS, MAB and KCW performed data analysis and interpreted results from sequencing and metabolomic data. ASD, IL, and MS handled mice and sample collection. RAD performed DNA and RNA extractions and quality control. RMF and IL wrote scripts to link ASVs and MAGs. YK, MB, and VHW were responsible for metabolomic analysis and LC-MS/MS sample prep and data collection. ASD and BMA handled *Salmonella* aspects of the project.

**Ikaia Leleiwi:** Conceptualization, Methodology, Software, Validation, Formal analysis, Investigation, Data Curation, Writing – Original Draft, Writing – Review & Editing, Visualization **Katherine Kokkinias:** Conceptualization, Validation, Formal analysis, Investigation, Data Curation, Writing – Review & Editing **Yongseok Kim:** Methodology, Validation, Formal analysis, Investigation, Data Curation, Writing – Original Draft **Maryam Baniasad:** Methodology, Validation, Formal analysis, Investigation, Data Curation, Writing – Review & Editing **Michael Shaffer:** Software, Investigation **Anice Sabag-Daigle:** Methodology, Validation, Investigation **Rebecca A. Daly:** Methodology, Software, Validation, Formal analysis, Investigation, Data Curation, Supervision, Project administration **Rory M. Flynn:** Software **Vicki H. Wysocki:** Conceptualization, Resources, Supervision, Project administration, Funding acquisition **Brian M. M. Ahmer:** Conceptualization, Resources, Writing – Review & Editing, Supervision, Project administration, Funding acquisition **Mikayla A. Borton:** Conceptualization, Software, Validation, Writing – Review & Editing, Supervision, Project administration, Funding acquisition **Kelly C. Wrighton:** Conceptualization, Resources, Writing – Review & Editing, Supervision, Project administration, Funding acquisition.

---

**Fig S1:**
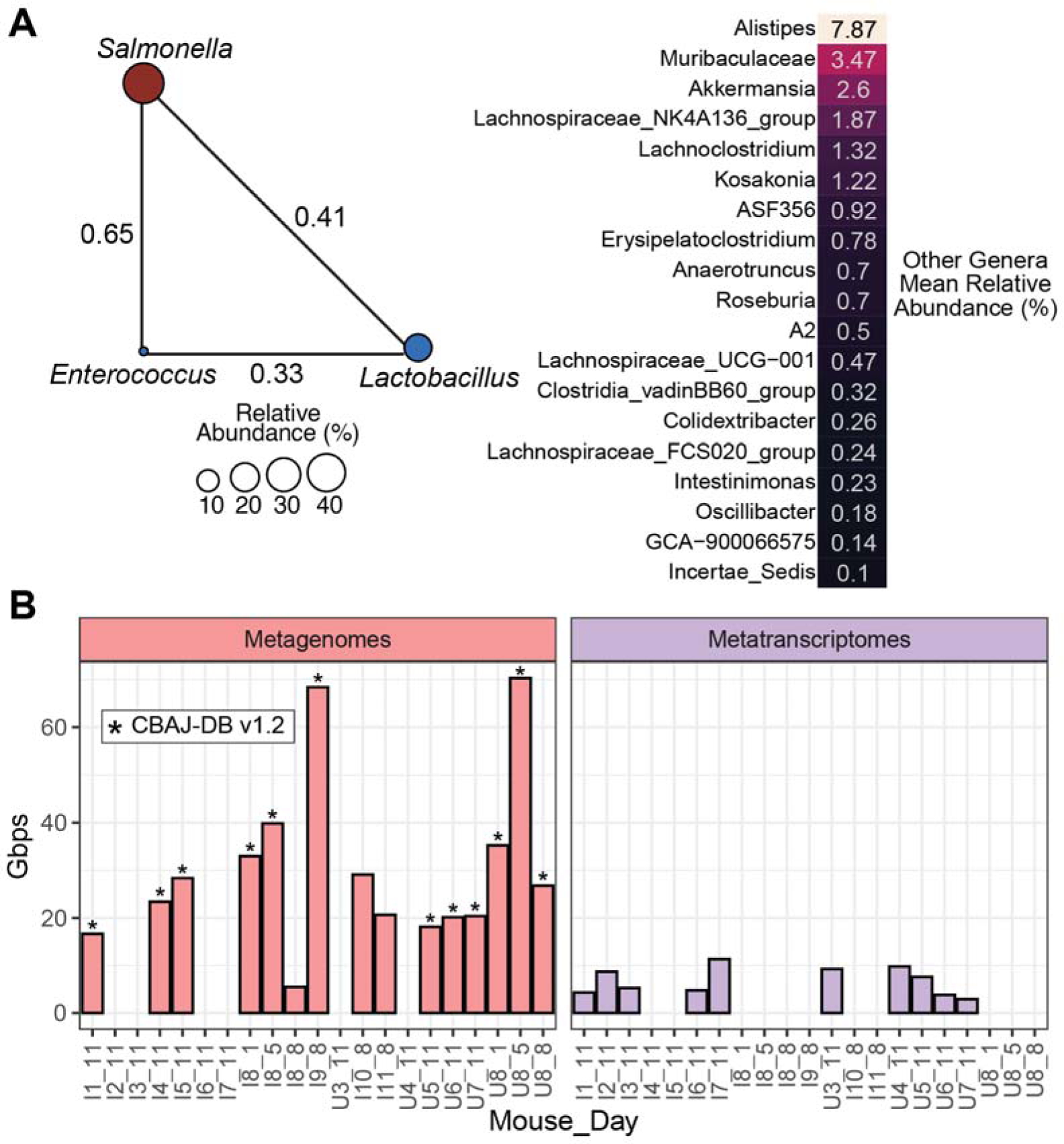
*Salmonella* correlation with *Bacilli* genera in later stages of infection and multi-omics sequencing effort. **A** Positive significant (p<0.05) Spearman correlations between community genera and *Salmonella*. Edges are the interaction Spearman correlation coefficient. Taxa listed on the right are all genera, and their mean relative abundances (16S rRNA), that displayed at least one significant positive correlation to another bacteria in later stage (days 10-12) infected samples. **B** Metagenomic and metatranscriptomic sequencing depth. Bars with asterisks indicate metagenomes used to produce the CBAJ-DB v1.2.

**Fig. S2:**
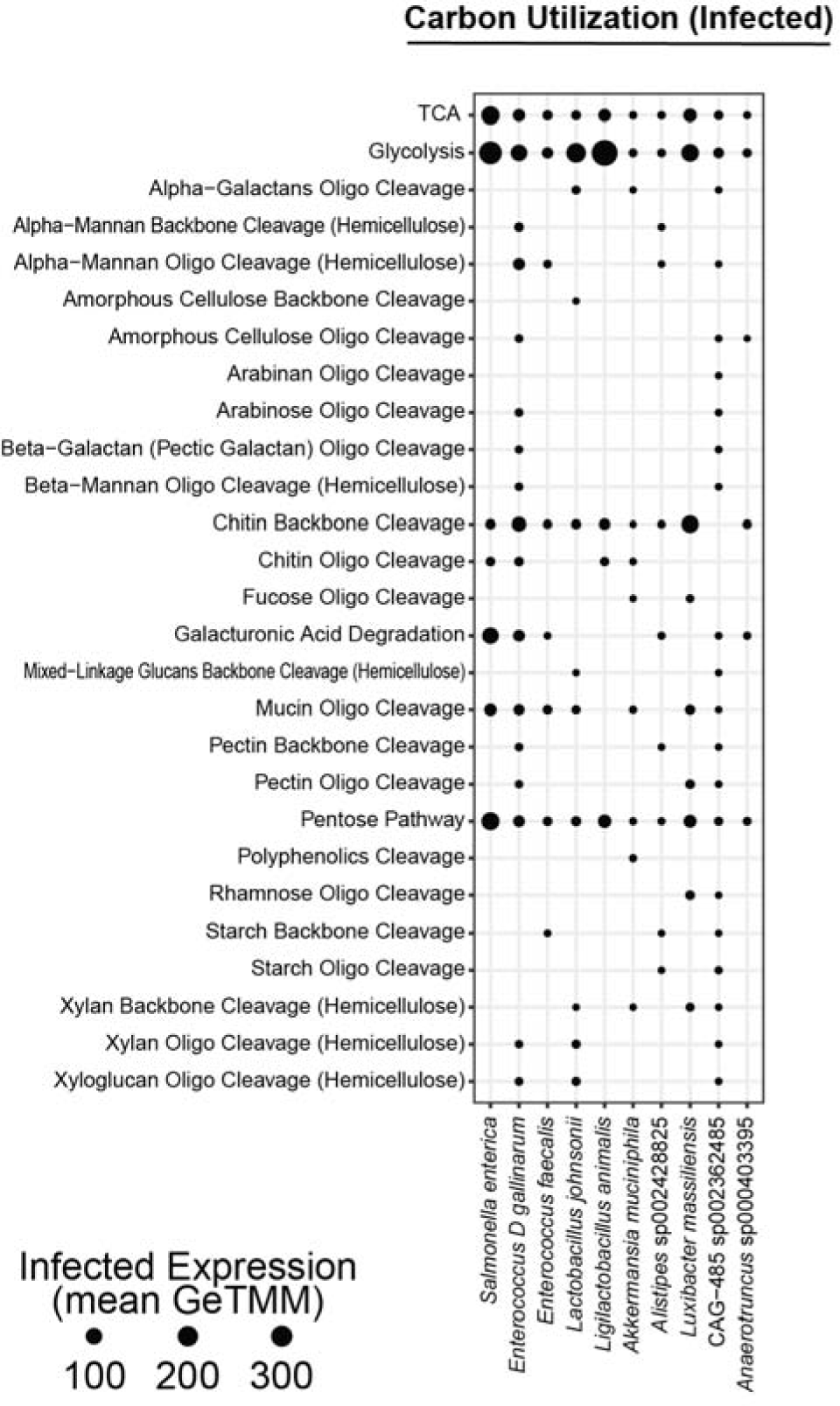
Carbon utilization of important microbiota in the infected gut. Carbon utilization of the most active and relatively abundant bacteria in the infected gut as determined by mapped metatranscriptomic reads to individual MAGs. Utilization categories were assigned by DRAM v1.4. Backbone cleavage denotes the presence of an endo-cleaving glycoside hydrolase and oligo cleavage denotes the presence of a exo-cleaving glycoside hydrolase.

**Fig S4:**
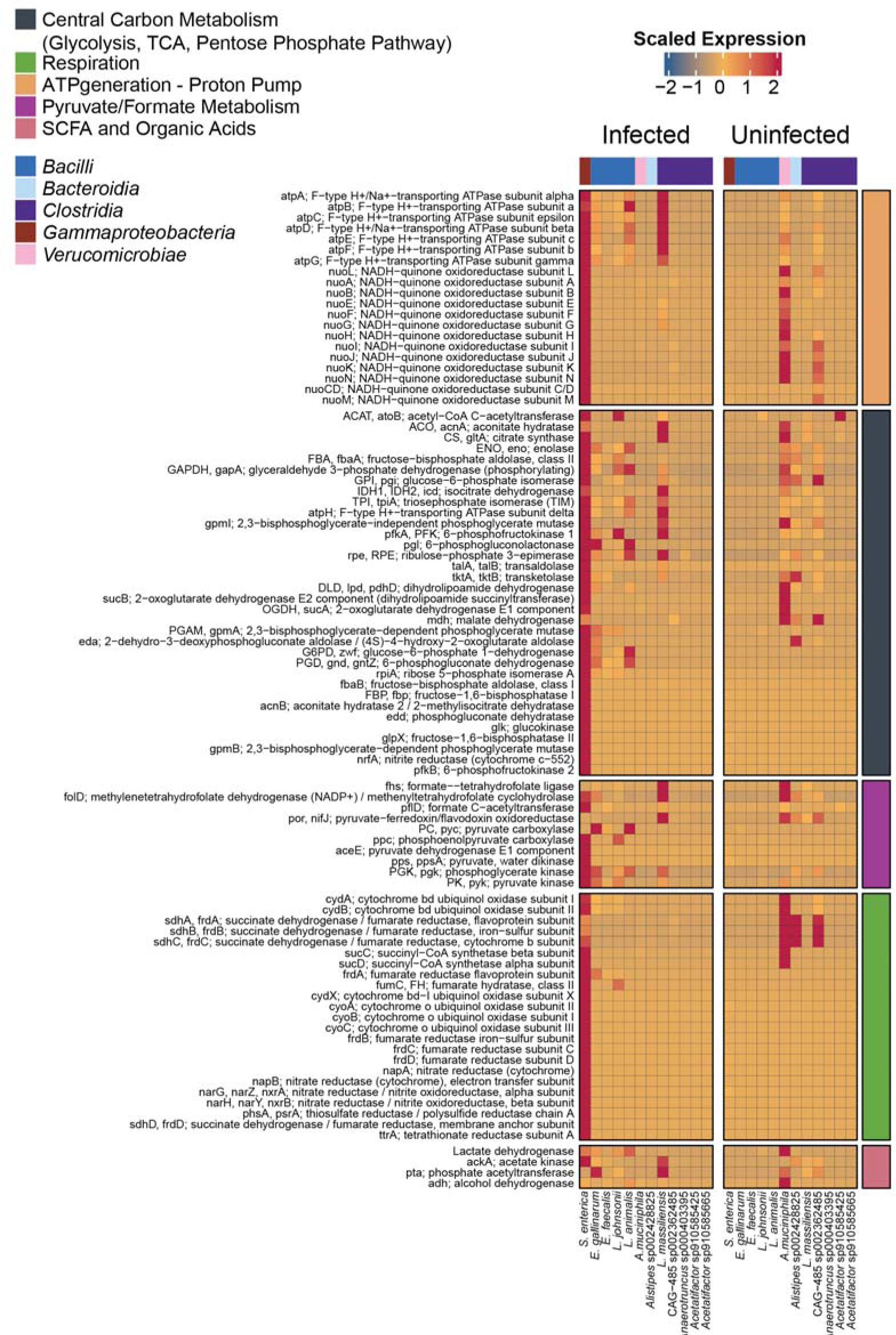
Differential gene expression of genes for energy production show altered metabolic strategies between treatments in prominent infected community members. Heatmap of actively expressed genes for energy production, including respiration, with select substrate utilization genes that are turned on during infection and are also significantly differentially expressed between treatments. Cell values are GeTMM scaled totals of all genes linked with a particular gene description (y-axis) averaged across samples within a treatment and expressed by an individual taxon (x-axis). Colored boxes on the right indicated gene class.

